# Leveraging *Drosophila* Models to Explore AI-generated Synthetic Peptide’s Potential in Boosting Honeybee Health and Resilience

**DOI:** 10.1101/2024.12.24.630224

**Authors:** Yanying Sun, Wenjie Jia, Yanan Wei, Xinyue Zhou, Ran Dong, Jihyeon Lee, Jeong Kyu Bang, Fangyong Ning, Woo Jae Kim

## Abstract

The integration of artificial intelligence (AI) and machine learning (ML) in peptide design has revolutionized the development of antimicrobial peptides (AMPs), which are essential components of innate immunity. In this study, we identified a novel synthetic peptide, PAN4 (GAYTFKIRRK), through genetic screening of AI-generated candidates in *Drosophila melanogaster*. PAN4 demonstrated robust antimicrobial activity, stress tolerance, and antitumor effects, significantly enhancing survival rates following bacterial infections and improving locomotor behaviors without adversely affecting lifespan. Furthermore, PAN4 expression in intestinal stem cells completely suppressed Ras^V12^-induced tumor progression, indicating its potential role in cancer prevention. The peptide also mitigated gut barrier dysfunction associated with sleep deprivation and reduced inflammation in a dextran sulfate sodium (DSS)-induced colitis model. Mechanistically, PAN4’s antimicrobial activity was linked to its interaction with specific peptidoglycan recognition proteins (PGRPs), particularly PGRP-SC1a, while the Tak1-mediated immune signaling pathway was found to be non-essential for its efficacy. PAN4 showed promising effects on honeybee health, enhancing survival rates under bacterial stress. Furthermore, PAN4 expression demonstrated significant anti-tumor activity in the *Drosophila* gut tumor model. Our findings suggest that PAN4 serves as a versatile agent with significant implications for enhancing immune responses and combating diseases in honeybee populations, paving the way for future applications in agriculture and medicine.

## INTRODUCTION

The integration of artificial intelligence (AI) and machine learning (ML) into peptide design represents a transformative shift in the field of biotechnology, particularly in the development of antimicrobial peptides (AMPs). AMPs are crucial components of the innate immune response, exhibiting broad-spectrum antimicrobial activity against bacteria, fungi, and viruses ^1,2^. However, the traditional methods of peptide discovery and optimization are often labor-intensive and time-consuming. The advent of AI-driven approaches has significantly enhanced the efficiency and efficacy of peptide design, enabling researchers to explore vast sequence spaces and identify novel peptides with desired therapeutic properties ^3–6^.

AI methodologies, particularly deep learning techniques, have been increasingly employed to streamline the peptide design process. These approaches utilize large datasets of known peptides to train models that can predict the biological activity, stability, and binding affinity of new peptide sequences. For instance, deep generative models (DGMs) such as variational autoencoders (VAEs) and generative adversarial networks (GANs) have been utilized to generate novel peptide sequences by learning from existing data distributions. This allows for the efficient exploration of sequence variations that might not be intuitively considered by human designers ^7–9^.

*Drosophila melanogaster*, commonly known as the fruit fly, serves as an exceptional model organism for testing synthetic peptides, particularly those with antimicrobial properties. Several factors contribute to its suitability, including its genetic tractability, rapid life cycle, and the availability of advanced genetic tools ^10–13^. *Drosophila* is also amenable to a variety of behavioral assays that can be used to assess the bioactivity of peptides. Its small size and rapid reproduction make it feasible to conduct large-scale experiments with minimal resources ^14^.

The fruit fly’s innate immune system shares significant similarities with that of higher organisms, making it an effective model for studying AMPs. *Drosophila* possesses a robust immune response characterized by the activation of antimicrobial peptide genes in response to pathogen infection ^11,15,16^. This feature allows for direct testing of peptide efficacy against bacterial infections in vivo ^17,18^. In summary, *Drosophila melanogaster* is a powerful model organism for testing synthetic peptides due to its genetic tractability, rapid life cycle, and sophisticated behavioral assays.

In this study, we report the identification of a highly effective AMP, designated ’PAN4’ (GAYTFKIRRK), which was discovered through genetic screening of AI-generated synthetic peptide candidates. PAN4 exhibits robust antimicrobial activity, as well as significant stress resistance and antitumor properties.

Furthermore, the application of PAN4 in honeybees has demonstrated beneficial effects on their health. Our findings suggest that the *Drosophila* platform serves as an effective screening system for evaluating a diverse array of peptides, with potential applications aimed at enhancing the health and resilience of honeybee populations.

## RESULTS

### Limited Efficacy of AI-Generated AMP Candidates in the *in vivo Drosophila* Model

To evaluate the potential of *Drosophila* as a testing platform for AI-generated synthetic AMPs, we initially assessed clinically validated peptides recognized as strong candidates for effective AMPs when expressed genetically ^3,4^. To evaluate the functionality of genetically encoded synthetic AMPs in the *Drosophila* model, we constructed *UAS-AMPs* that include a previously characterized signal peptide at their N-termini ^19^. Among the peptides tested, none exhibited antimicrobial activity against *Pseudomonas aeruginosa*, a gram-negative pathogen that commonly afflicts humans, insects, and plants ^18,20^ (Fig. S1a-e). Recent advancements in machine learning techniques, particularly artificial neural networks, have enhanced computer-based approaches to AMP design ^3,8^.Utilizing various computational strategies for AMP design, we selected a set of candidates derived from AI-driven methodologies ^21–27^. However, similar to the clinically proven peptides, none of the AI-designed peptides demonstrated antimicrobial activity against *P. aeruginosa* (Fig. S1f-n). These findings indicate that the effective design of AMPs for specific applications is a complex challenge that necessitates careful consideration of both host and target specificity ^28,29^.

Next, we turn our attention to an innovative approach utilizing Generative Adversarial Networks (GANs) for the design of AMPs. GANs are a powerful class of machine learning models that consist of two neural networks—the generator and the discriminator—engaged in a competitive process. The generator creates new data instances, while the discriminator evaluates these instances against real data, providing feedback that enhances the generator’s output. This adversarial training allows GANs to produce high-quality synthetic data that closely resembles the training dataset, making them particularly effective for generating complex biological sequences such as peptides ^8,30–34^.

The results from a recent study demonstrated that several peptides designed using the GAN framework, referred to as PandoraGAN ^35^, exhibited promising antiviral activity *in vitro*, underscoring the potential of AI-driven methodologies to enhance peptide discovery. This research illustrates the effectiveness of GANs in addressing the limitations encountered by traditional AMP design strategies and presents a novel approach for developing effective antiviral agents. Among the five peptides generated (Fig. S2a), we identified PAN4 (Fig. 1a) and PAN5 as exhibiting strong antimicrobial activity (Fig. S2b-e). Consequently, we selected PAN4 as a promising candidate for further testing as a beneficial AMP for honeybee health (Fig. 1b-c). We genetically encoding PAN4 construct comprising a trypsin signal sequence for secretion, the bioactive PAN4 peptide, and a hydrophilic linker with a c-myc tag for structural flexibility and detection (Fig.1d).

**Fig 1.**
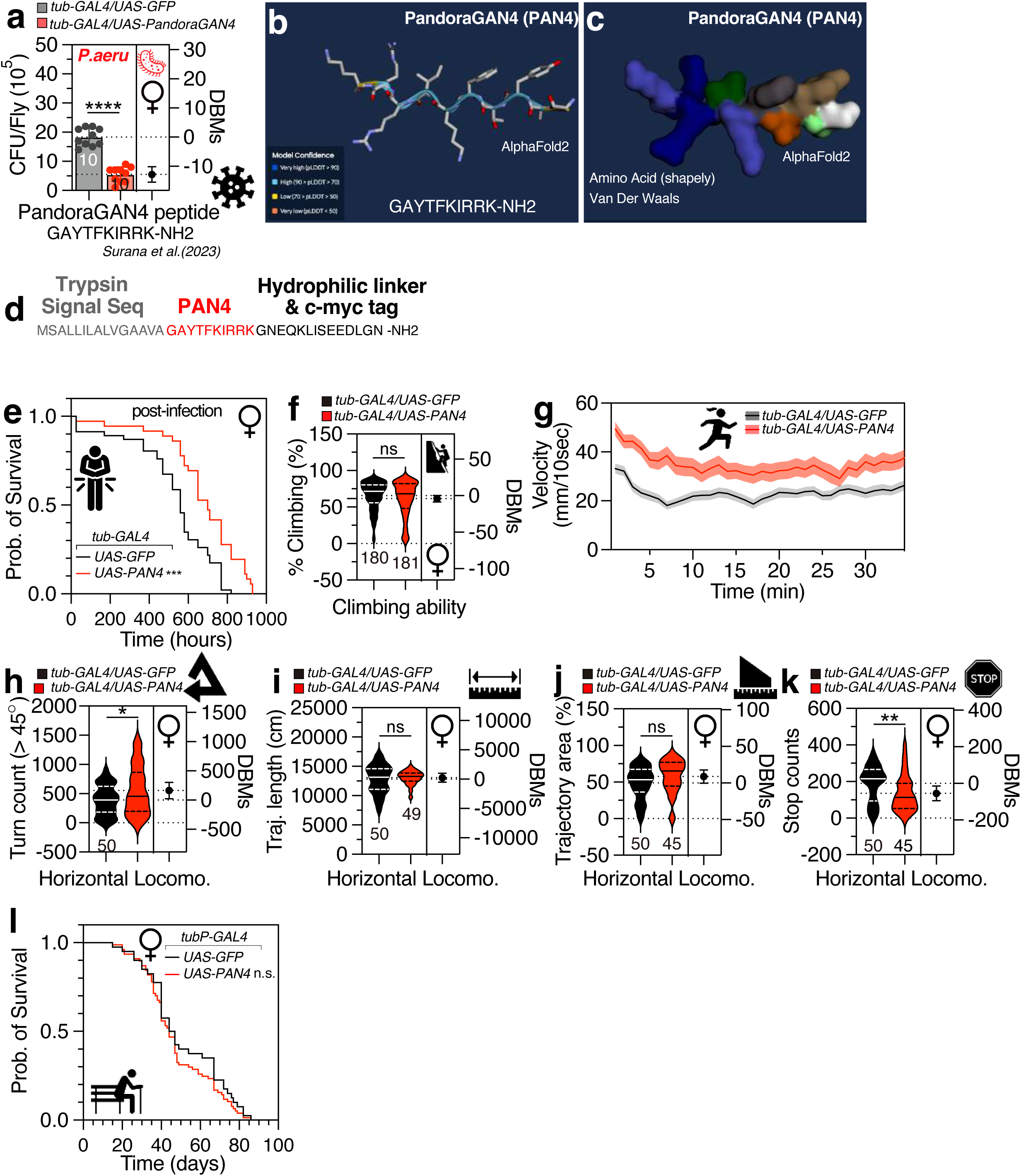
Antimicrobial activity of PandoraGAN4 (PAN4), structural and sequence analysis of its amino acids, survival and locomotion of flies expressing PAN4. (A) Infectious dose of female flies expressing PandoraGAN4 (PAN4) by *tub-GAL4*. (B-C) Predicted structure and amino acid sequence features of PAN4 generated by AlphaFold2. (D) The amino acid sequence of PAN4, including trypsin signal sequence (grey), PAN4 sequence (red), hydrophilic linker sequence (black). (E) The survival curve of female PAN4 flies post infection by *P. aeruginosa* culture. (F) The climbing ability, (G) velocity, (H) turn count, (I) trajectory length, (J) trajectory area, and (K) stop counts of female PAN4 flies. (L) The survival curve of female flies expressing PAN4 via *tub-GAL4*. The mean value and standard error are labeled within the dot plot (black lines). DBMs represent the ’difference between means’ for the evaluation of estimation statistics (See **METHODS**). Asterisks represent significant differences, as revealed by the unpaired Student’s t test, and ns represents non-significant differences (*\*p<0.05, **p<0.01, ***p< 0.001, ****p< 0.0001*).

### Genetic expression of PAN4 beneficially changes gut environment and enhances Overall Health in *Drosophila*

Genetically expressed PAN4 significantly enhanced survival rates in *Drosophila* following infection (Fig. 1e). While broad expression of PAN4 did not affect the climbing ability of the flies (Fig. 1f), it did improve locomotor behaviors, including forward velocity (Fig. 1g) and rotation frequency (Fig. 1h). However, trajectory length (Fig. 1i) and trajectory percentage (Fig. 1j) remained unchanged, while the number of stops decreased (Fig. 1k). The lifespan of flies expressing PAN4 was comparable to that of the genetic control group (Fig. 1l). Collectively, these data indicate that widespread expression of the secreted form of PAN4 in *Drosophila* confers potent antimicrobial activity, modifies locomotor behaviors, and does not adversely affect fly lifespan.

To assess the efficacy of genetically expressed PAN4 against various bacterial infections, we infected fruit flies with the Gram-positive pathogen *Enterococcus faecalis*, commonly associated with intestinal infections, and found that PAN4 expression did not inhibit *E. faecalis* infection (Fig. S3a). Furthermore, PAN4 expression was ineffective in blocking infection by *Escherichia coli* (Fig. S3b). Additionally, PAN4 expression did not mitigate the infection caused by *Lactobacillus plantarum*, a bacterium frequently found in fermented foods and utilized as a probiotic (Fig. S3c). These findings indicate that PAN4 may exhibit selective inhibition of specific bacterial species.

To determine the specific tissues in which PAN4 exerts its antimicrobial effects, we employed various GAL4 drivers to express PAN4 in targeted tissues (Fig. 2a-l). Our findings revealed that expressing PAN4 in either enteroendocrine cells (EEs) (Fig. 2j-k) or intestinal stem cells (ISCs) (Fig. 2l) was sufficient to replicate the effects observed with *tub-GAL4* expression. This suggests that EEs and ISCs play crucial roles in PAN4’s ability to eliminate harmful bacteria within the fly gut.

**Fig 2.**
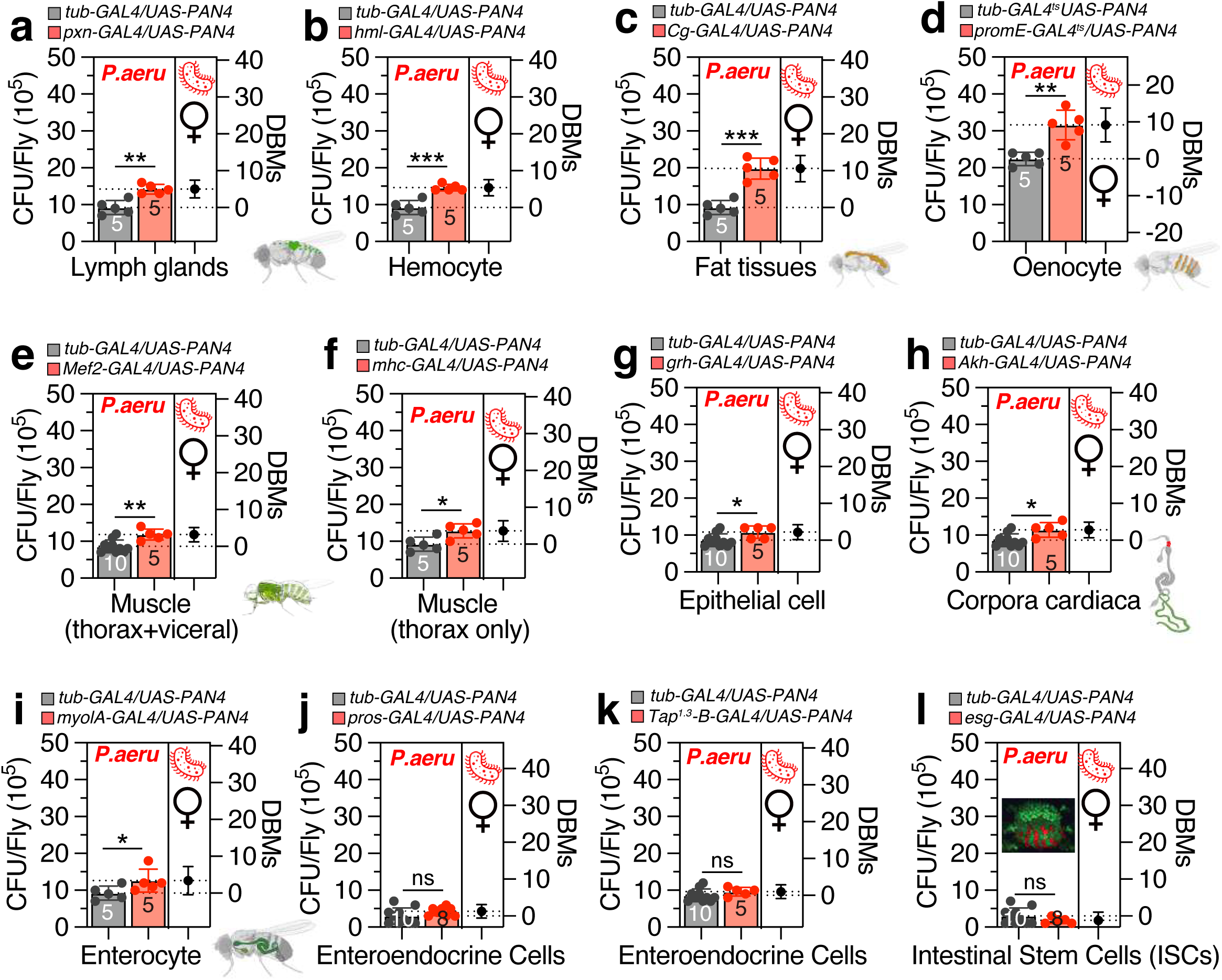
Tissue screening of antimicrobial effects in PAN4 expressing female flies. (A-L) Infectious dose between female flies expressing PAN4 via *tub-GAL4* and *Pxn-Gal4* (A), *Hml-GAL4* (B), *Cg-GAL4* (C), *promE-GAL4* (D), *Mef2-GAL4* (E), *Mhc-GAL4* (F), *grh-GAL4* (G), *Akh-GAL4* (H), *MyolA-GAL4* (I), *pros-GAL4* (J), *Tap^1.3^-B-GAL4* (K)*, esg-GAL4* (L), following a 12-hour orally feeding of *P. aeruginosa* culture.

Notably, the expression of PAN4 in ISCs resulted in a significant reduction in stem cell numbers in both the R2 and R4 regions of the fly gut (Fig. 3a-b), indicating that PAN4 may influence stem cell proliferation (Fig. 3c-d). However, PAN4 expression markedly enhanced mitochondrial activity in the most regions of the gut (Fig. 3e-h and Fig. S3d-f), suggesting that PAN4 stimulates mitochondrial biogenesis. Additionally, expression of PAN4 in EEs led to a significant decrease in midgut pH, rendering it more acidic, as demonstrated by phenol-red dye absorption (Fig. 3i). Interestingly, the genetic expression of PAN4 in EEs also resulted in a dramatic increase in calcium levels in the R3 region of the gut (Fig. 3j-l). However, the increase in calcium levels was not observed in the ovaries (Fig. S3g-i), indicating that the calcium elevation induced by PAN4 is confined to the midgut. Therefore, the potent antimicrobial activity of PAN4 appears to be attributed to enhanced mitochondrial activity, acidification of midgut pH, and activation of calcium signaling in enteroendocrine cells.

**Fig 3.**
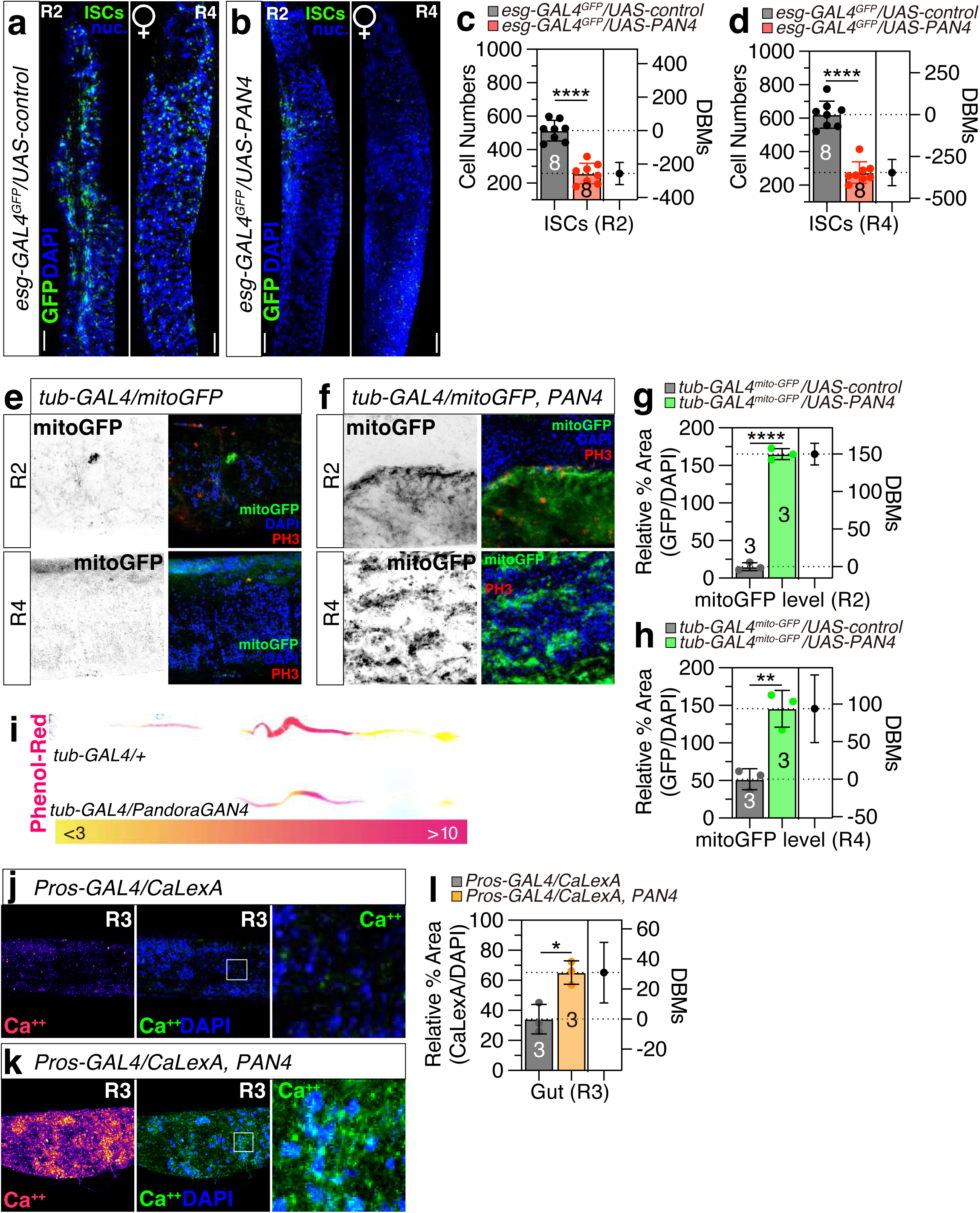
PAN4 suppresses intestinal stem cell proliferation and alters mitochondrial and calcium signaling in *Drosophila*. (A-D) Representative images and quantification of intestinal stem cells (ISCs) by *esg-GAL4* with *UAS-GFP* flies expressing *UAS-control* or *UAS-PAN4*. Fly guts were immunostained with anti-GFP (green), and DAPI (blue). Scalebar 200µm. (E-H) Mitochondrial activity visualized by mitoGFP and phosphorylated histone H3 (PH3) staining by *tub-GAL4*-driven *UAS-control* or *UAS-PAN4* expression and quantification of GFP fluorescence in R2 (G) and R4 region (H). (I) pH changes assessed by phenol-red staining in normal and PAN4 expressing conditions via *tub-GAL4* driver, flies were fed by phenol red dye which changes from yellow at pH<3, an acidic region, to brighter red at pH>10, an alkaline region. (J-L) Calcium signaling alterations assessed by CaLexA reporter in for *Pros-GAL4* together with *lexAop-mCD8GFP; UAS-CaLexA*, *lexAop-CD2-GFP* of control (J) and PAN4 expressed (K) female flies at R3 region of gut, and were immunostained with anti-GFP (green), and DAPI (blue), calcium signal was quantified as relative percentage area normalized by DAPI signal (L). Scalebar 200µm. Asterisks represent significant differences, as revealed by the unpaired Student’s t test, and ns represents non-significant differences (*\*p<0.05, **p<0.01, ***p< 0.001, ****p< 0.0001*).

### PAN4 Expression Potentially Alleviates Stress-Induced Pathological Responses While Modestly Affecting Sleep and Foraging Behaviors

Given that PAN4 expression in the gut enhances overall fly health, we investigated whether it could also mitigate stress responses. Sleep deprivation can lead to increased mortality due to the accumulation of reactive oxygen species (ROS) in the gut, with gut neuropeptides playing a role in mediating energy depletion caused by sleep loss in *Drosophila* ^36,37^. Sleep deprivation disrupts the integrity of the intestinal epithelial barrier, a phenomenon that reflects an evolutionarily conserved aspect of aging and is associated with changes in metabolic and inflammatory biomarkers ^38,39^. Our results indicate that PAN4 expression can alleviate gut barrier dysfunction induced by sleep deprivation ^40^ (Fig. 4a-b and Fig. S4a). Therefore, the presence of PAN4 in the gastrointestinal tract of *Drosophila* may act as a protective mechanism against ROS-mediated stress resulting from sleep loss.

**Fig 4.**
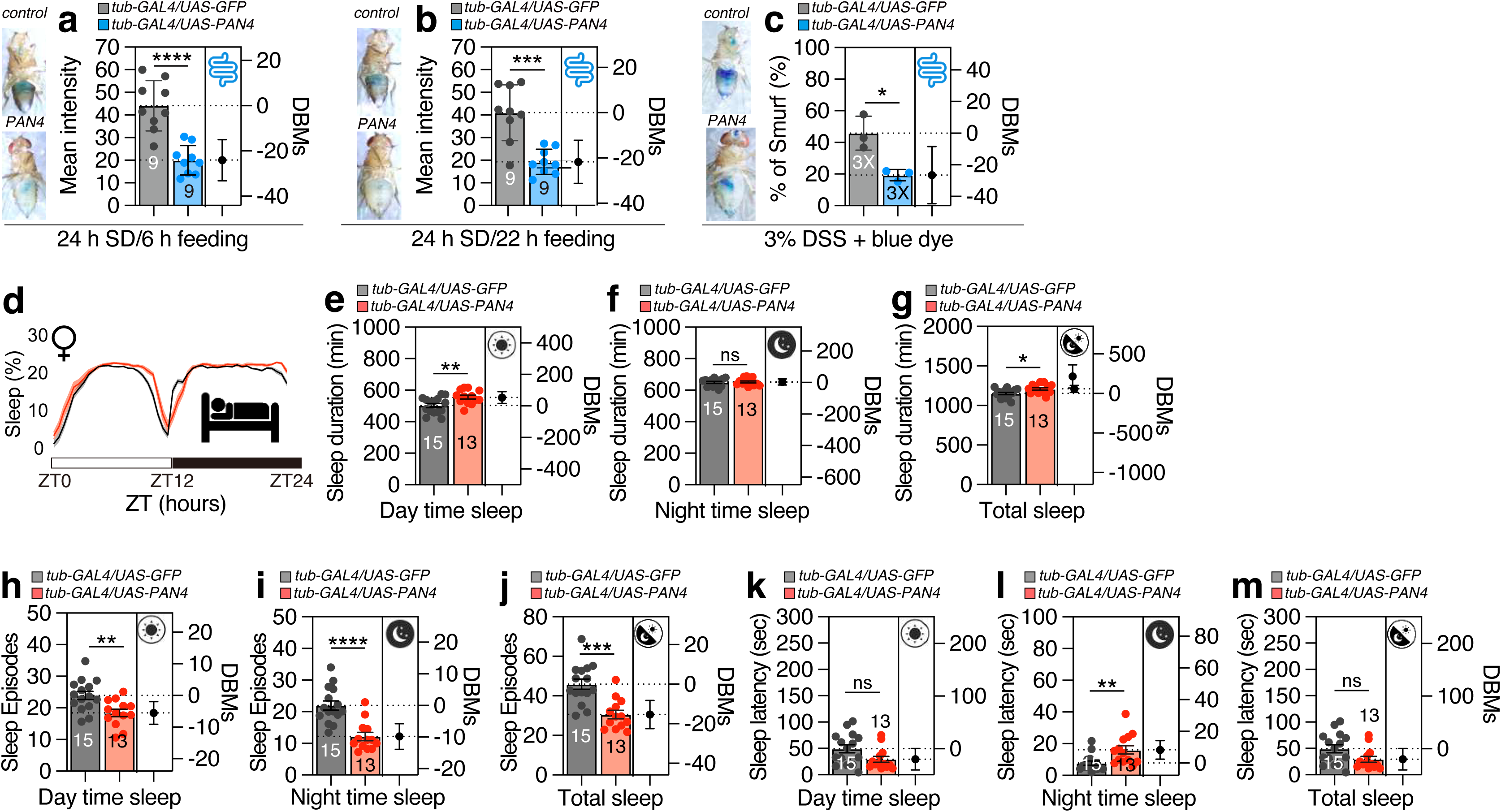
Stress-induced Smurf phenotypes and sleep patterns of pan-expressed PAN4 fly. (A-B) The mean intensity of Smurf assay flies expressing PAN4 compared with controls under sleep deprivation. The experiment was repeated three times and one representative figure was shown of each condition. The mean intensity of Smurf assay flies expressing PAN4 compared with controls after 24 hours sleep deprivation and 6 hours (A) and 22 hours (B) dye feeding. The experiment was repeated three times and one representative figure was shown of each condition. (C) The mean intensity of Smurf assay flies expressing PAN4 treated with a 3% DSS feeding with blue dye. The experiment was repeated three times and one representative figure was shown of each condition. (D-M) Sleep profiles (average proportion of time spent sleeping in consecutive 30-min segments during a 24-h LD cycle) and quantification of sleep parameters of female flies expressing PAN4 expression by *tub-GAL4* driver. Quantification of sleep eduration for PAN4 expression flies in (E) to (G). Quantification of sleep episodes for PAN4 expression flies in (H) to (J). Quantification of sleep latency for PAN4 expressing flies in (K) to (M). Asterisks represent significant differences, as revealed by the unpaired Student’s t test, and ns represents non-significant differences (*\*p<0.05, **p<0.01, ***p< 0.001, ****p< 0.0001*).

Additionally, administration of dextran sulfate sodium (DSS) induces mucosal injury in the adult *Drosophila* gastrointestinal tract, adversely affecting viability ^41,42^. The DSS-induced intestinal inflammation model has become widely recognized as an effective experimental framework for studying colitis and inflammatory bowel disease (IBD) ^43,44^. Our findings demonstrate that PAN4 expression can mitigate DSS-induced intestinal inflammation (Fig. 4c), suggesting that PAN4 acts as a potent inhibitor of DSS-mediated epithelial damage.

Given that PAN4 can mitigate ROS stress induced by sleep deprivation, we investigated its effects on sleep behavior. Expression of PAN4 resulted in increased daytime sleep duration and a reduction in the number of sleep episodes, while also extending nighttime sleep latency (Fig. 4d-m). These findings suggest that PAN4 positively influences sleep behavior. Additionally, we examined whether PAN4 expression affects the foraging behavior of flies and discovered that it significantly enhanced the foraging activity of well-fed flies (Fig. S4b-e). This indicates that PAN4 expression not only promotes increased foraging activity but also improves sleep quality, contributing to the overall health and well-being of the flies.

### Specific Peptidoglycan Recognition Proteins (PGRPs) are Essential for the Antimicrobial Activity of PAN4

Peptidoglycan recognition proteins (PGRPs) play a critical role in detecting peptidoglycan, a key component of bacterial cell walls. The interaction between PGRPs and peptidoglycan triggers a signaling cascade that activates immune responses, including the Toll and immune deficiency (Imd) pathways, leading to the production of AMPs mediated by the transcription factor Relish (Rel) ^10,12^. In *Drosophila*, the PGRP family includes various members such as PGRP-L (long forms), PGRP-S (short forms), as well as transmembrane, intracytoplasmic, and secreted proteins. These proteins are distributed across multiple tissues and cell types, including hemocytes, epithelial cells, and fat body cells, and are essential for the proper functioning of the immune system ^45–47^ (Fig. 5a).

**Fig 5.**
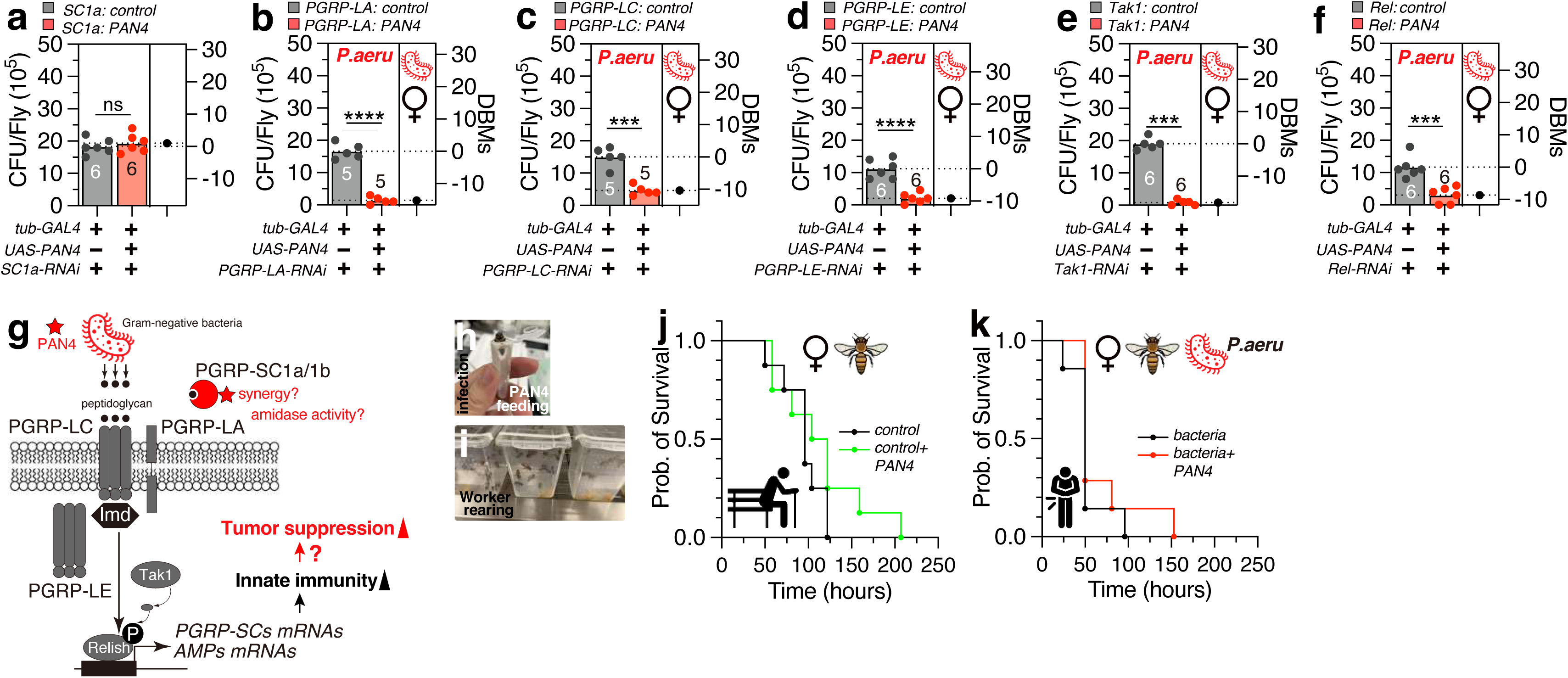
Schematic depicting the role of PAN4 in immune pathway gene screening, its function in antimicrobial peptides (AMP), application in honeybees and its potential anti-tumor effects. (A-F) Infectious dose of controls and female flies expressing PAN4 via *tub-GAL4*, with a knockdown of *PGRP-SC1a* (A), *PGRP-LA* (B), *PGRP-LC* (C), *PGRP-LE* (D), *Tak1* (E), *Rel* (F), following a 12-hour orally feeding of *P. aeruginosa* culture. (G) The diagram of PAN4 involved in PGRP family related immune pathway. (H-I) Honeybee PAN4 feeding (H) and infection assay application. Worker bee rearing (I). (J) The survival curve of honeybees in control (black) compared with high concentration PAN4 (green) administration. (K) The survival curve of honeybees infected orally by P. *aeruginosa*, control (black) compared with high concentration PAN4 (red) administration. (L-N) Immunostaining indicating PAN4 expression in the midgut with control and expressing *Ras^v12^via esg^ts^-GAL4*. Transgenes were induced with *esg^ts^* by incubating at 29[°C for indicated durations. The cells manipulated by *esg^ts^*are marked and stained with GFP (green), anti-PH3 with RFP (Red) and nuclei are stained with DAPI (blue). Scalebar 200µm. (O) Quantification of PH3-positive cells per adult midgut of the indicated genotypes. Asterisks represent significant differences, as revealed by the unpaired Student’s t test, and ns represents non-significant differences (*\*p<0.05, **p<0.01, ***p< 0.001, ****p< 0.0001*).

Knockdown of PGRP-SC1a abolished the antimicrobial effect of PAN4 (Fig. 5a), indicating that PAN4’s antimicrobial activity relies on the amidase activity of PGRP-SC1a to combat bacterial infections ^46,48,49^. In contrast, knockdown of PGRP-LA, LC, and LE did not affect PAN4’s antimicrobial efficacy (Fig. 5b-d), suggesting that the antimicrobial action of PAN4 does not depend on these PGRPs.

In *Drosophila*, Tak1 (Transforming Growth Factor-β Activated Kinase 1) is a key player in the Imd signaling pathway, which is vital for the innate immune response against Gram-negative bacterial infections ^50,51^. Relish serves as a crucial downstream component of the IMD pathway that regulates antibacterial responses ^12^. Notably, knockdown of either Tak1 or Rel did not reduce the antimicrobial activity of PAN4 (Fig. 5e-f), suggesting that the Tak1-mediated kinase cascade and Rel-mediated transcriptional regulation are not necessary for PAN4’s antimicrobial properties (Fig. 5g).

Honeybees, particularly *Apis mellifera* and *Apis cerana*, are essential for the pollination of numerous crops and play a critical role in maintaining ecosystem stability ^52^. Unfortunately, honeybee populations are currently declining due to various threats, including pests, pathogens, genetic bottlenecks, and environmental challenges. This decline poses significant risks to both agriculture and the environment ^53^.

AMPs and other peptides produced by honeybees are integral to their innate immune defense mechanisms. These peptides effectively target and eliminate a wide array of pathogens, making them promising candidates for novel genetic strategies aimed at enhancing honeybee immunity ^54^. However, the molecular mechanisms governing the function and regulation of these peptides remain largely unexplored.

To evaluate the applicability of PAN4’s protective effects observed in the *Drosophila* model to honeybee physiology, we conducted a series of experiments involving the dietary administration of PAN4 to honeybees (Fig. 5j-k). The introduction of PAN4 into their diet resulted in significantly adverse effects on honeybee viability. Notably, PAN4 supplementation led to an increase in the lifespan of worker honeybees isolated from their colonies (Fig. 5j), suggesting that PAN4 may confer survival benefits. Furthermore, PAN4 supplementation improved the survival rate of worker honeybees exposed to a high bacterial load (Fig. 5k). These findings indicate that PAN4 provides protective effects against bacterial infections in the honeybee gut, similar to those observed in the *Drosophila* model.

Recent studies have highlighted a significant relationship between immune signaling pathways and cancer progression. For instance, chronic activation of the Imd pathway can lead to dysregulation of epithelial homeostasis, promoting conditions conducive to tumor growth. Specifically, PGRP-SCs has been shown to regulate ISC proliferation and maintain gut immune homeostasis, its downregulation is associated with increased ISC hyperproliferation and epithelial dysplasia, which are hallmarks of tumorigenesis ^55,56^. This suggests that PGRPs not only play a protective role against pathogens but may also influence tumor dynamics ^57–60^.

In this context, we hypothesized that PAN4 could influence the immune response through PGRPs while also affecting tumor growth in our *Drosophila* model. The expression of PAN4 in intestinal stem cells (ISCs) significantly inhibits tumor progression induced by both Ras^V12^ and Yki^3SA^ (Fig. 6a-b). Upon closer examination, we found that PAN4 expression nearly eliminates Ras^V12^-expressing malignant tumors in the midgut of *Drosophila*, restoring it to a normal gut state (Fig. 6c-f). Notably, the survival of PAN4-expressing flies with Ras^V12^ tumors is nearly twice as long as that of the control group with only Ras^V12^ tumors (Fig. 6g). These findings strongly suggest that the AI-generated synthetic peptide PAN4, validated in our fly model, exhibits potent anti-tumor activity against both Ras^V12^-and Yki^S3A^-induced malignant gut tumors.

**Fig 6.**
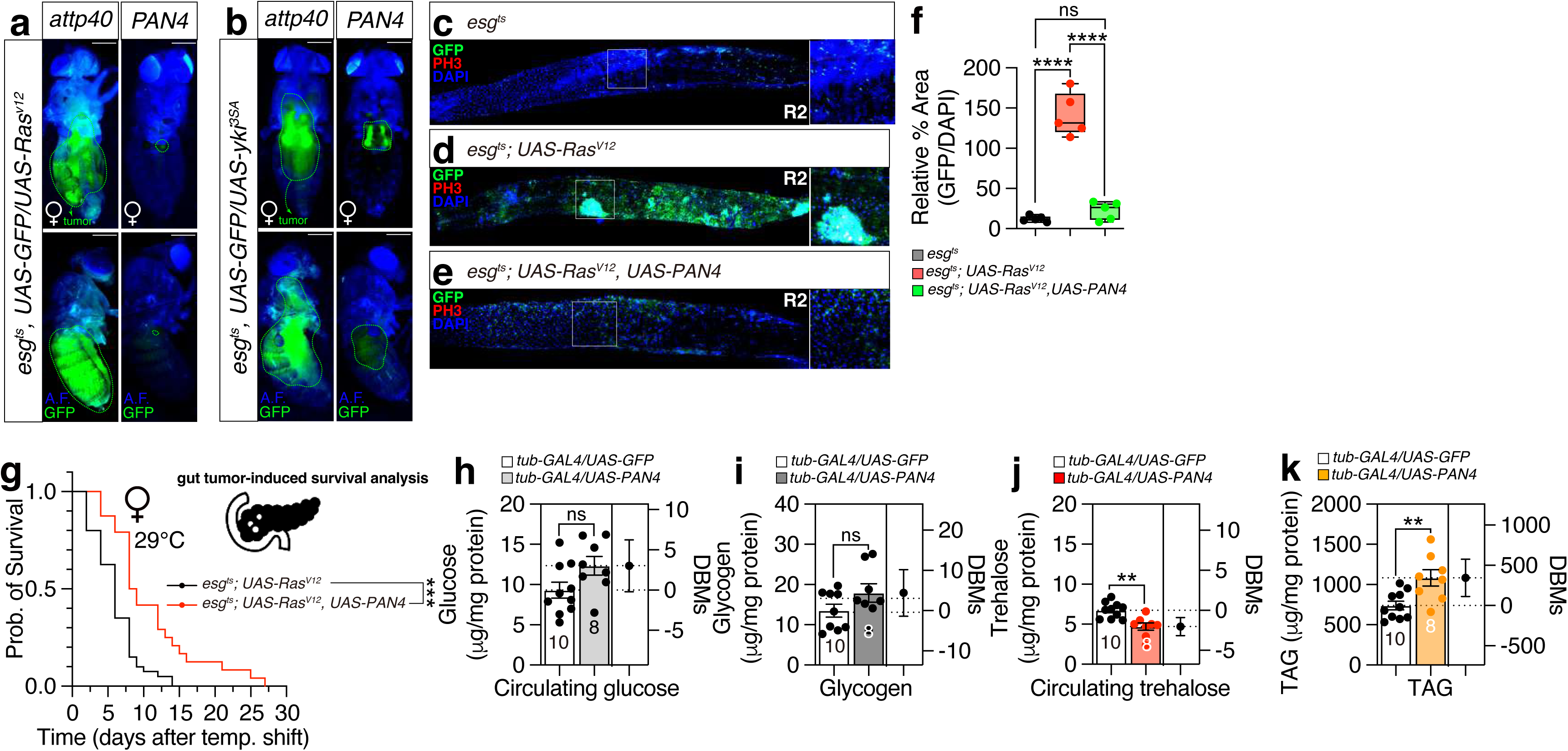
PAN4 suppresses malignant gut tumors and modulates metabolic homeostasis in *Drosophila*. (A-B) Expression of PAN4 significantly inhibits tumor progression induced by *Ras^V12^* or *Yki^3SA^* in *Drosophila*. A.F.means autofluorescence. (C-F) Immunostaining indicating PAN4 expression in the midgut R2 region with control and expressing *Ras^v12^via esg^ts^-GAL4*. Transgenes were induced with *esg^ts^* by incubating at 29[°C for indicated durations. The cells manipulated by *esg^ts^* are marked and stained with GFP (green), anti-PH3 with RFP (Red) and nuclei are stained with DAPI (blue). Scalebar 200µm. (G) Survival curve of PAN4-expressing flies with *Ras^V12^* tumors. Transgenes were induced with *esg^ts^* by incubating at 29[°C for indicated durations. (H-K) PAN4 expression modulates metabolic homeostasis in *Drosophila*.PAN4 expression normalizes glucose and glycogen levels while reducing trehalose and increasing triacylglycerol (TAG) levels. Asterisks represent significant differences, as revealed by the unpaired Student’s t test, and ns represents non-significant differences (*\*p<0.05, **p<0.01, ***p< 0.001, ****p< 0.0001*).

It is known that malignant tumors in *Drosophila* disrupt insulin signaling ^61^, leading to increased gluconeogenesis ^62^. Flies expressing PAN4 displayed glucose and glycogen levels comparable to those of control flies (Fig. 6h-i), but they exhibited decreased circulating trehalose levels (Fig. 6j) and increased triacylglycerol (TAG) levels (Fig. 6k). In *Drosophila*, glucose levels fluctuate more dynamically than trehalose levels in the circulating hemolymph in response to dietary changes, with trehalose metabolism acting as a buffer for circulating glucose levels ^63^. TAG serves as a crucial caloric source for energy homeostasis in animals, and its turnover is vital for the metabolism of structural and signaling lipids. These neutral lipids also play significant roles in development and disease ^64^. Interestingly, studies have demonstrated that tumors induced by Yki in flies exhibit elevated levels of trehalose while showing a decrease in TAG levels ^65^. The observed decrease in trehalose levels alongside an increase in TAG levels due to PAN4 expression may help explain its tumor-suppressive effects.

Our results suggest that the antimicrobial properties of PAN4 may extend beyond pathogen defense, potentially providing insights into its role in inhibiting tumor development through modulation of the Imd pathway, PGRP activity, and metabolic processes related to gluconeogenesis.

## DISCUSSION

In this study, we explored the multifaceted benefits of the AI-engineered synthetic peptide PAN4 (GAYTFKIRRK) in *Drosophila*, demonstrating its potent antimicrobial, stress-tolerant, and antitumor properties. Through genetic screening and AI-driven design methodologies, PAN4 was identified as an effective antimicrobial peptide that enhances survival rates following bacterial infections while positively influencing locomotor behaviors and overall health. Our findings revealed that PAN4 expression in ISCs completely suppresses Ras^V12^-induced tumor progression, indicating its potential role in cancer prevention. Furthermore, PAN4 mitigates stress responses associated with sleep deprivation and DSS-induced gut inflammation, highlighting its protective effects against ROS and intestinal barrier dysfunction. The antimicrobial activity of PAN4 is closely linked to the function of specific PGRPs, particularly PGRP-SC1a, while the Tak1-mediated Imd signaling pathway appears non-essential for its efficacy. Additionally, we assessed the applicability of PAN4 in honeybees, revealing its potential to enhance honeybee health by improving survival rates under bacterial stress.

PAN4 exerts its antimicrobial and antitumor effects through several molecular mechanisms, primarily by modulating the immune response in *Drosophila*. The peptide’s interaction with PGRPs initiates a signaling cascade that activates the Imd pathway, leading to the production of AMPs via the transcription factor Rel (Fig. 5g) ^66^. Specifically, our findings indicate that PAN4’s antimicrobial activity is dependent on the amidase activity of PGRP-SC1a, which is essential for combating bacterial infections. Additionally, PAN4 enhances mitochondrial activity and promotes gut health by acidifying midgut pH and activating calcium signaling in enteroendocrine cells, thereby contributing to its protective effects against both bacterial infections and tumorigenesis ^67^. These mechanisms highlight the potential of PAN4 as a multifaceted agent that not only targets pathogens but also influences key cellular processes involved in maintaining gut homeostasis and preventing tumor progression.

The successful application of AI-generated peptides like PAN4 in *Drosophila* raises exciting possibilities for their use in other model organisms and practical applications in agriculture and medicine. The ability to engineer peptides with specific antimicrobial properties could be invaluable in developing new treatments for diseases affecting crops and livestock, thereby enhancing food security and agricultural sustainability. Furthermore, as honeybee populations face significant challenges from pathogens and environmental stressors, peptides like PAN4 could be integrated into strategies aimed at improving bee health and resilience against infections. This demonstrates the versatility of AI-driven peptide design in addressing a wide array of biological challenges.

To fully elucidate the mechanisms by which PAN4 interacts with immune pathways and its broader physiological effects, several avenues for future research are warranted. Detailed studies are needed to investigate the specific molecular interactions between PAN4 and PGRPs, as well as the downstream signaling events activated by these interactions. Additionally, exploring the effects of PAN4 on other physiological processes, such as metabolic regulation and stress response pathways, will provide a more comprehensive understanding of its potential benefits. Investigating the long-term effects of PAN4 supplementation on health outcomes in both *Drosophila* and honeybees will also be crucial for assessing its viability as a therapeutic agent.

## MATERIALS AND METHODS

### AlphaFold Structure Prediction

To predict the three-dimensional structure of the synthetic peptide PAN4 (GAYTFKIRRK), we utilized the AlphaFold2 protein structure prediction tool, which has been recognized for its high accuracy in modeling protein structures. The AlphaFold2 algorithm is based on deep learning techniques and has demonstrated significant advancements in protein folding predictions, as detailed by Jumper et al. (2021) in *Nature* ^68,69^. The predicted structure was visualized and analyzed using molecular visualization software to assess the conformation and potential functional sites of PAN4.

### Generation of synthetic PAN4 peptide

The peptide were prepared by Fmoc based solid phase peptide synthesis (SPPS) methods using Rink amide resin with an initial loading of 0.44 mmol/g from amino acids containing fluorenylmethoxycarbonyl (Fmoc) protecting group, all the amino acids were purchased from Novabiochem. Full peptide sequence obtained from microwave-assisted peptide synthesizer (CEM, Liberty blue model). The resin was swollen in N, N’-Dimethylformamide (DMF) for 10 min prior for synthesis. For the sequence extension, we have utilized the Fmoc protected amino acids (5.0 equiv) Oxyma pure (5.0 equiv). The synthetic method was carried out by gradually raising the temperature from 60 °C to 75°C within 2 min, the entire reaction condition kept under nitrogen atmosphere. In addition, each deprotection process accompanied in the same method using 20% piperidine/DMF solvent system. For double cyclization, the side chains at Cys 1 and Cys 11 positions were protected with Trt groups, and mmt groups were used at Cys 3 and Cys 15 positions. After the reaction was completed, the mmt group was removed by treating three times with 2% TFA/DCM at 1-minute intervals. Subsequently, for cyclization on the resin, 300µL of 0.06M iodine/DMF solution was added to the resin swelled in 3 mL DMF and allowed to vortexing for 15 min. Later, washed the resin by ascorbic acid 1M DMF solution for removing iodine residues. After successful processing, the resin was washed three times each with DMF, DCM, and methanol, dried well, and the peptides were cleaved with 95/2.5/2.5 (TFA/TIS/H2O) for 3 h. The product was obtained by precipitation in ethyl ether.

Afterwards, the peptide from which all protecting groups, including trt, were removed, was purified through HPLC, followed by lyophilization, then, dissolved in DMSO solvent for the second cyclization. DIPEA (10%) was added and reacted for overnight. Finally purification was carried out in HPLC to achieve the double-cyclized peptide.

### Fly Stocks and Husbandry

*Drosophila melanogaster* was raised on cornmeal-yeast medium at similar densities to yield adults with similar body sizes. Flies were kept in 12 h light: 12 h dark cycles (LD) at 25 [(ZT 0 is the beginning of the light phase, ZT12 beginning of the dark phase). To reduce the variation from genetic background, all flies were backcrossed for at least 3 generations to CS strain. All mutants and transgenic lines used here and their sources were as follows: *w^1118^*(Vienna Drosophila Resource Center, 60000), *w^1118^;UAS-mito-HA-GFP/CyO* (Bloomington Stock Center, 8442), *w^1118^;Cg-GAL4* (Bloomington Stock Center, 7011), *;;tub-GAL4/TM3*, *;;Hml-GAL4*, ;*MyolA-Gal4,UAS-GFP/CyO*, *w*; esg-Gal4,UAS-GFP/CyO, ;Pxn-GAL4(II)/CyO, ;; pros-GAL4/TM3, ; esg^ts^/CyO, UAS-GFP; UAS-Ras^v12^/Ubx, esg^ts^/CyO, UAS-GFP; UAS-Yki^3SA^/Ubx* (Theses lines were kindly provided by Dr. Lihua Jin, Northeast Forestry University, Harbin, China), *;promE-GAL4,Tub-GAL80ts(II)/CyO* (Korea Drosophila Resource Center, 2161), *w*;; mef2-GAL4* (Bloomington Stock Center, 50742), *;;Mhc-GAL4(III)/TM3* (Bloomington Stock Center, 55133), *W^1118^; grh-GAL4* (Bloomington Stock Center, 65637), *;; Akh-GAL4* (Bloomington Stock Center, 25684), *;; tap^1.3^-B-GaL4* (The line was kindly provided by Dr. Zheng Guo, Huazhong University, Wuhan, China), *;LexAop-CD8GFP(II);UAS-CaLexA,LexAop-CD2-GFP/TM6B,Tb* (Korea Drosophila Resource Center, 1234), *y1 sc* v1 sev21; PGRP-SC1a-RNAi* (Bloomington Stock Center, 57184), *;PGRP-LA-RNAi* (Vienna Drosophila Resource Center, 102277), *y^1^*,*sc*,v^1^,sev^21^;; UAS-PGRP-LC-RNAi* (Bloomington Stock Center, 33383), *y^1^*,*sc*,v^1^,sev^21^;UAS-PGRP-LE-RNAi* (Bloomington Stock Center, 60038), *y^1^ v^1^; UAS-Tak1-RNAi* (Bloomington Stock Center, 53377), *y^1^,sc*,v^1^,sev^21^;; Rel-RNAi* (Bloomington Stock Center, 33661).

### Molecular Cloning and Injection for Transgenic Fly Generation

To generate the *UAS-AMPs* used in this study, peptide cDNAs were chemically synthesized with optimal *Drosophila* codon usage and with an optimal *Drosophila* Kozak translation initiation site upstream of the start methionine (CAAA). PAN4 Encoded peptides is GAYTFKIRRK. A trypsin signal peptide (NSELINSLLSLPKNMNDA) for secretion was incorporated at the N-terminus of the PAN4 sequence, followed by a hydrophilic linker sequence (GNEQKLISEEDLGN) that includes an embedded c-Myc epitope tag, as outlined by Choi et al. (2009) ^19^. The cDNA was cloned into the pUAS-attB vector; For generation of transgenic *Drosophila*, Vectors was injected into the embryos of flies. The genetic construct was inserted into the attp40 site on chromosome II to generate transgenic flies using established techniques, a service conducted by Qidong Fungene Biotechnology Co., Ltd. (http://www.fungene.tech/).

### Lifespan Assay and Statistical Analysis

For lifespan analysis, we used conventional procedure as we described before ^70^. Briefly, 50 flies were aged by sex before being raised in typical 12 h light: 12 h dark cycles at either 25°C for each experimental objective. The number of dead flies was recorded every two to three days. Every three to four days, the surviving flies were transferred to fresh vials.

### Climbing assay

For climbing assay, we modified the conventional RING assay ^71^. In brief, 40-50 20-day-aged flies were placed in an empty vial and were tapped to the bottom of the tube. After tapping of flies, we recorded 10 seconds of video clip. This experiment was done five times with 5-minute intervals. With recorded video files, we captured the position of flies 10 seconds after tapping the vial. This captured image file was then loaded in ImageJ to perform particle analysis. For quantifying the location of flies inside a vial, we used the “analyze particles” function of ImageJ ^72^. The position of pixels was normalized by height of vial then only the particles above the midline (4 cm) of vial were counted.

### Locomotion assay

To detect and quantify the activity of flies, we have developed the Fly Trajectory Dynamics Tracking (FlyTrDT) software. This is an open-source, custom-written Python program that utilizes the free OpenCV machine vision library and the Python Qt library. The FlyTrDT software simultaneously records the trajectory information of each fly and calculates various indicators of the group at a certain period. For each frame acquired, the moving fly is segmented using the binarization function from the OpenCV library. Subsequently, a Gaussian blur and morphological closing and opening operations were performed on the extracted foreground pixels to consolidate detected features and reducing false positives and negatives. Finally, the extraction of fly outlines was achieved using the contour detection algorithm in the OpenCV library.

### Bacteria culture

*P. aeruginosa* (ATCC 27853) culture, 10 mL of Luria-Bertani (LB) broth was inoculated with 100 µL of a frozen bacterial stock at 37 °C. The main procedure was modified based on previous study^73^. Shake at 150 rpm overnight and grow this subculture in a 1 L conical flask for another night. Pour equal volumes of this subculture across 500 mL centrifuge tubes and spin the subculture at 2,500 x g for 15 min at 4 °C to pellet the bacteria. Remove the supernatant and resuspend the final bacteria pellet in 5% sucrose water solution. Check the OD and adjust to the desired infectious dose (OD600 = 25 in this study).

### Bacterial infection assay

The main procedure was modified based on previous study and described as follows ^73^. Flies were starved for 4 h before exposure to bacteria by transferring the flies to empty vials. Place a disc of filter paper on top of food and pipette 100 µL of bacterial culture directly onto the filter disc. For control infections in this study, bacterial culture was replaced with PBS. 5 flies were transferred to the sample tube and leave for 12 h infection exposure. To confirm oral infection, first surface-sterilize the flies immediately after bacterial exposure, by placing them in 100 µL of 70% ethanol for 20–30s. Remove the ethanol and add 100 µL of triple distilled water for 20–30s before removing the water. Add 100 µL of 1x PBS and homogenize the fly. Transfer the homogenate to the top row of a 96-well plate and add 90 µL of 1x PBS to every well below. Serially dilute this sample to distinguish a range of CFU values. Take 10 µL of the homogenate in the top well and add this to the well below. Repeat this step with the second well, transferring 10 µL to the third well, and so on, for as many serial dilutions as required. Plate the serial dilutions on an LB nutrient agar plate in 2 µL droplets, to ensure all droplets remain discrete. Incubate the LB Agar plates overnight at 37 °C and count visible CFUs. Calculate the number of CFUs per fly by counting the number of colonies present at the serial dilution where 0–40 CFUs are clearly visible. Then check the colony numbers in 10^-5^.

### Immunostaining

After 5 days of eclosion, the *Drosophila* gut was taken from adult flies and fixed in 4% formaldehyde at room temperature for 30 minutes. The sample was than washed three times (5 minutes each) in 1% PBT and then blocked in 5% normal goat serum for 30 minutes. Subsequently, the sample was incubated overnight at 4 [with primary antibodies in 1% PBT, followed by the addition of fluorophore-conjugated secondary antibodies for one hour at room temperature. For DAPI staining, the gut was incubated in DAPI containing 1% PBT for 10 minutes at room temperature, followed by three times wash. Finally, the gut was mounted on plates with an antifade mounting solution (Solarbio) for imaging purposes. Samples were imaged with Zeiss LSM880. Antibodies were used at the following dilutions: Chicken anti-GFP (1:500, Invitrogen), rabbit anti-PH3 (1:200, Merck), Alexa-488 donkey anti-chicken (1:200, Jackson ImmunoResearch), Alexa-555 donkey anti-rat (1:200, Invitrogen), DAPI (1:1000, Invitrogen).

### Phenol red pH test

Flies were transferred to food containing 0.2% Phenol Red (Macklin, P6066) ^74^, and fed for at least 6 hours. Gut was dissected in PBS and imaged under microscope.

### Smurf assay

For Smurf assay, 3% (v/v) Food Blue No.1 aluminum lake (Aladdin, F336821) was mixed with normal food, and flies under different conditions were imaged on Olympus SZ61 microscope after a 6-hour feeding or 20-hour feeding^75^. For flies under sleep deprivation, a vortex machine (CHANGZHOU ENPEI INSTRUMENT MANUFACTURING CO., LTD., NY-5SX) was applied with a routine of 2-second 1500 rpm vortex following a one-minute rest^76^, flies were transferred in tubes containing normal food and went through a 24-hour sleep deprivation. For DSS feeding, 3% DSS (Coolaber, 9011-18-1) was mixed with blue dye food and flies were transferred and fed overnight ^77^.

### Single-fly sleep and circadian rhythm recording

96-well white Microfluor 2 plates (Fishier) with 400 μl of food (5% sucrose and 1% agar) were loaded with adult male flies (aged 3–5 days). Flies were entrained to the 12[h:12[h LD cycles for four days at 25[°C to record sleep behavior, then changed to constant darkness for 5-6 days to record circadian rhythms in the absence of light inputs. The fly movement was monitored using a camera at 10s intervals, and the data were then used by the sleep and circadian analysis program SCAMP to analyze sleep and circadian rhythm ^78–80^. It calculates activity by shifting the position of *Drosophila* every 10 seconds and calculates sleep using the standard definition (*Drosophila* is recorded as asleep if it remains motionless for at least 5 minutes).

### Honeybee infection assay

The European honeybees, *Apis mellifera*, in this study were kindly provided and maintained by Dr. Fangyong Ning (Northeast Agricultural University, Harbin, China). For oral administration, the procedure was modified based on previous study, and was described as follow ^81^. Bees were anesthetized on ice and placed in 1.5 ml tubes with the assistance of paper tapes for fixation. Bees were recovered from anesthesia at room temperature and fed 10 µl of each sample. For control groups, bees were administrated with 1 mol/L sucrose and for bacterial infection group, bees were fed with bacteria pellet resuspended by 1 mol/L sucrose. As for PAN4 administration, PAN4 powder (Chemstan, 24345-16-2) was dissolved into 1 mol/L sucrose at a high concentration of 0.01mg/mL, and 100 µL of high concentration PAN4 was transferred to 1.5 ml tube, adding 900 µL 1 mol/L sucrose creating a medium concentration. Repeat this step to create low concentration PAN4, and PAN4 at various concentrations were applied to honeybees using a micropipette. Treated bees were maintained in a plastic box and fed with small honeycomb at 25°C. Honeybees were judged to be dead and recorded if no motion was detected in any body parts.

### Tumor GFP intensity measurement from confocal images

GFP was quantitated from full projections of images acquired using confocal microscopy. GFP intensity in gray scale from regions of interest (ROIs) covering the entire gut was acquired using the Leica-LSM proprietary software. GFP intensity was normalized to the area of each ROI or DAPI signal. Student’s *t*-test was performed using GraphPad Prism to look for statistical significance in GFP variation ^82^

### Tumor quantification and quantification of phospho-histone H3 (PH3)-positive cells

Protocol to analyze *Drosophila* intestinal tumor cellular heterogeneity using immunofluorescence imaging and nuclear size quantification ^83^ To assess the number of dividing cells, midguts were dissected and stained with DAPI and an anti-PH3 antibody. PH3-positive nuclei were quantified across the entire midgut.

### Nutrient-level assays

Flies were collected at 7 days of age, with 100 flies used for analysis. Groups of 10 flies were placed into 1.5 mL Eppendorf tubes and snap-frozen at -80°C. The flies were homogenized in 300 μL PBST (PBS + 0.5% Tween-20) using a homogenizer. Protein concentration was quantified by mixing 2 μL of homogenate with 100 μL of Bradford reagent, measuring absorbance at 595 nm, and generating a standard curve for normalization. Samples were then heat-inactivated at 70°C for 5 minutes, centrifuged at maximum speed for 3 seconds, and the supernatant was transferred to a new tube. For triacylglyceride (TAG) measurement, 2 μL of supernatant was mixed with 4 μL triacylglyceride reagent (Sigma, T2449) and 6 μL PBST, incubated at 37°C for 30 minutes, followed by the addition of 20 μL free glycerol reagent (Sigma, F6428) and incubation at 37°C until color development (30–60 minutes), with absorbance measured at 540 nm and quantified using a glycerol standard curve. For glycogen measurement, 1 μL homogenate was mixed with 0.2 μL amyloglucosidase (Sigma, A7420) diluted to 5 mg/mL and 9.8 μL PBST, incubated at 37°C for 30 minutes, followed by the addition of 25 μL glucose oxidase (GO) reagent (Sigma, GAGO20) and incubation at 37°C for color development, with absorbance measured at 540 nm. Free glucose was measured by mixing 2 μL homogenate with 25 μL GO reagent without amyloglucosidase, incubating at 37°C, and measuring absorbance at 540 nm. Trehalose measurement involved incubating 1 μL supernatant with 0.5 μL trehalase (Sigma, T8778) and 9.5 μL PBST overnight at 37°C, followed by the addition of 25 μL GO reagent, further incubation at 37°C, and absorbance measurement at 540 nm. Free glucose values were subtracted from total glucose + glycogen and trehalose + glucose values to determine glycogen and trehalose levels, respectively. All values were normalized to protein content for standardization.

## STATISTICAL ANALYSIS

All analysis was done in GraphPad (Prism9). When dataset passed test for normal distribution (Kolmogorov-Smirnov tests, p > 0.05), we used two-sided Student’s t tests. The mean ± standard error (s.e.m) (*\*\*\*\* = p < 0.0001, *** = p < 0.001, ** = p < 0.01, * = p < 0.05*). All analysis was done in GraphPad (Prism). Individual tests and significance are detailed in figure or figure legends.

Besides traditional *t*-test for statistical analysis, we added estimation statistics for all analysis in two group comparing graphs. In short, ‘estimation statistics’ is a simple framework that—while avoiding the pitfalls of significance testing—uses familiar statistical concepts: means, mean differences, and error bars. More importantly, it focuses on the effect size of one’s experiment/intervention, as opposed to significance testing ^84^. In comparison to typical NHST plots, estimation graphics have the following five significant advantages such as (1) avoid false dichotomy, (2) display all observed values (3) visualize estimate precision (4) show mean difference distribution. And most importantly (5) by focusing attention on an effect size, the difference diagram encourages quantitative reasoning about the system under study ^85^. Thus, we conducted a reanalysis of all our two group data sets using both standard *t* tests and estimate statistics. In 2019, the Society for Neuroscience journal eNeuro instituted a policy recommending the use of estimation graphics as the preferred method for data presentation ^86^.

## DATA AND CODE AVAILABILITY

Strains are available upon request. The authors affirm that all data necessary for confirming the conclusions of the article are present within the article, figures, and tables.

### ACKNOWLEDGEMENTS

We thank Lihua Jin (Northeast Forest University, China) and Zheng Guo (Huazhong University of Science and Technology, China) for sharing valuable fly strains. The genetic research conducted within this study was significantly enhanced by the comprehensive resources offered by the FlyBase website. We extend our gratitude to the FlyBase team for their diligent maintenance of this extensive Drosophila database ^87–91^. We also extend our gratitude to the fly resource centers, including BDSC, VDRC, KDRC, and Tsinghua Fly Center, for providing the fly strains used in this study. This research was financially supported by the Startup funds from the HIT Center for Life Science, awarded to WJK. It is important to note that the funding bodies played no role in the study design, data collection and analysis, decision to publish, or the preparation of the manuscript.

## ETHICS APPROVAL AND STATEMENT OF ANIMAL RESEARCH COMPLIANCE

All animal experiments reported in this manuscript were conducted in compliance with the ARRIVE guidelines and adhered to the U.K. Animals (Scientific Procedures) Act, 1986 and associated guidelines, EU Directive 2010/63/EU for animal experiments, or the National Research Council’s Guide for the Care and Use of Laboratory Animals ^92^.

## CONSENT TO PARTICIPATE

The research described in this paper does not involve any human participants. Therefore, no consent to participate was obtained.

## CONSENT FOR PUBLICATION

All authors have given consent for the publication of this work.

## DECLARATION OF INTEREST

The authors declare no competing interests.

## DECLARATION OF GENERATIVE AI AND AI-ASSISTED TECHNOLOGIES IN THE WRITING PROCESS

During the creation of this work, the author(s) utilized ChatGLM (https://chatglm.cn/) and KIMI (https://kimi.moonshot.cn/) to rephrase English sentences, verify English grammar, and detect plagiarism, as none of the authors of this paper are native English speakers. After using this tool/service, the author(s) reviewed and edited the content as needed and take(s) full responsibility for the content of the publication.

## AUTHOR CONTRIBUTIONS

**Conceptualization:** Woo Jae Kim.

**Data curation:** Yanying Sun, Wenjie Jia, Woo Jae Kim.

**Formal analysis:** Yanying Sun, Wenjie Jia, Woo Jae Kim.

**Funding acquisition:** Woo Jae Kim.

**Investigation:** Woo Jae Kim.

**Methodology:** Jihyeon Lee, Jeong Kyu Bang, Fangyong Ning, Woo Jae Kim.

**Project administration:** Woo Jae Kim.

**Resources:** Jihyeon Lee, Jeong Kyu Bang, Woo Jae Kim.

**Supervision:** Fangyong Ning, Woo Jae Kim.

**Validation:** Yanying Sun, Woo Jae Kim.

**Visualization:** Yanying Sun, Wenjie Jia, Woo Jae Kim.

**Writing – original draft:** Woo Jae Kim.

**Writing – review & editing:** Yanying Sun, Yanan Wei, Xinyue Zhou, Woo Jae Kim.

## SUPPLEMENTAL FIGURE TITLES AND LEGENDS

**Fig S1. Antimicrobial activity of clinical and AI-designed peptides against microbial pathogens.**

(A-N) Antimicrobial activity upon infection with P. *aeruginosa* in clinical tested peptides and f AI-designed peptides. Clinical tested peptides by feeding of *P. aeruginosa* in (A) to (E). AI-designed peptides following a 12-hour orally feeding of *P. aeruginosa* culture in (F) to (N).

**Fig S2. Sequences and antimicrobial activities of PandoraGAN-derived peptides.**

(A) Sequences of PandoraGAN-derived peptides.

(B-E) Antimicrobial activities of PandoraGAN peptides following a 12-hour orally feeding of *P. aeruginosa* culture.

**Fig S3. Infectious dose of flies expressing PAN4 encountering different bacteria and gut and ovary environment changes displayed.**

(A-C) Infectious dose of female flies expressing PAN4 by *tub-GAL4* following a 12-hour orally feeding of *E. faecalis* culture (A), *E. coli* culture (B), and *Lactobacillus plantarum* culture (C).

(D-F) The PH3-positive motisis cells and mitochondria (UAS-mitoGFP) in R3 region of female flies gut in normal (D) and PAN4 expressing conditions (E) by *tub-GAL4*, fly guts were immunostained with anti-PH3 (red), anti-GFP (green), and DAPI (blue).

Scalebar 200µm. GFP signal was quantified as relative percentage area normalized by DAPI signal (F).

(G-I) CalexA assay for *Pros-GAL4* together with *lexAop-mCD8GFP; UAS-CaLexA, lexAop-CD2-GFP* of control (G) and PAN4 expressed (H) female ovary, and were immunostained with anti-GFP (green), and DAPI (blue). Calcium signal was quantified as relative percentage area normalized by DAPI signal (I). Asterisks represent significant differences, as revealed by the unpaired Student’s t test, and ns represents non-significant differences (*\*p<0.05, **p<0.01, ***p< 0.001, ****p< 0.0001*).

**Fig S4. Smurf assay in normal condition and foraging activity in different condition.**

(A) The mean intensity of Smurf assay flies expressing PAN4 compared with control under normal condition. The experiment was repeated three times and one representative figure was shown of each condition.

(B-E) Foraging activity with control under fed and straved condition (B), expressing PAN4 with *tub-GAL4* under fed and starved (C), quantification of fed and starvation under control (D) and expressing PAN4 condition (E).

## Notes

### Competing Interest Statement

The authors have declared no competing interest.

