## Supplementary figures and images for "Leveraging *Drosophila* Models to Explore AI-generated Synthetic Peptide’s Potential in Boosting Honeybee Health and Resilience"

### Supplemental Figure.1

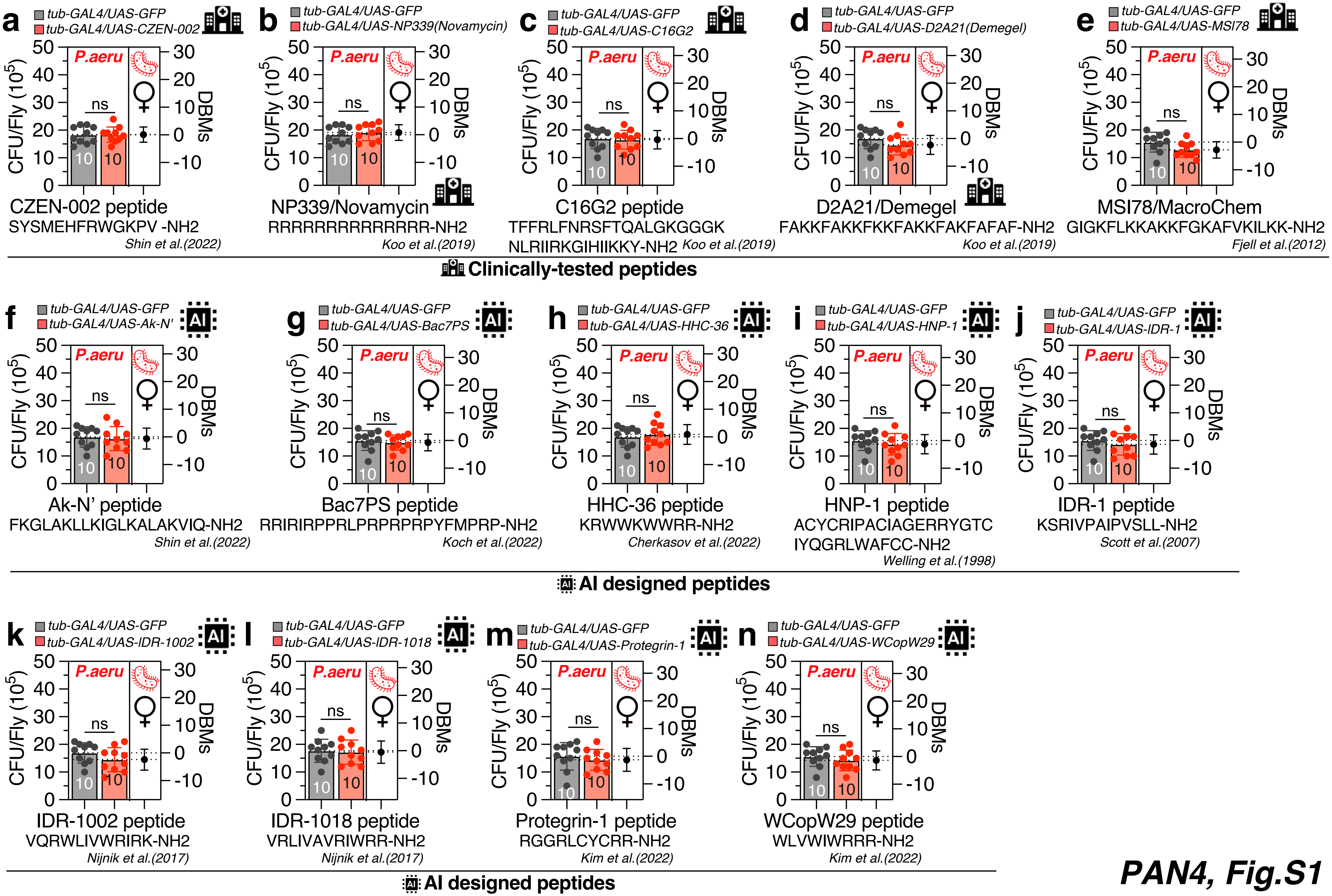

### Supplemental Figure.3

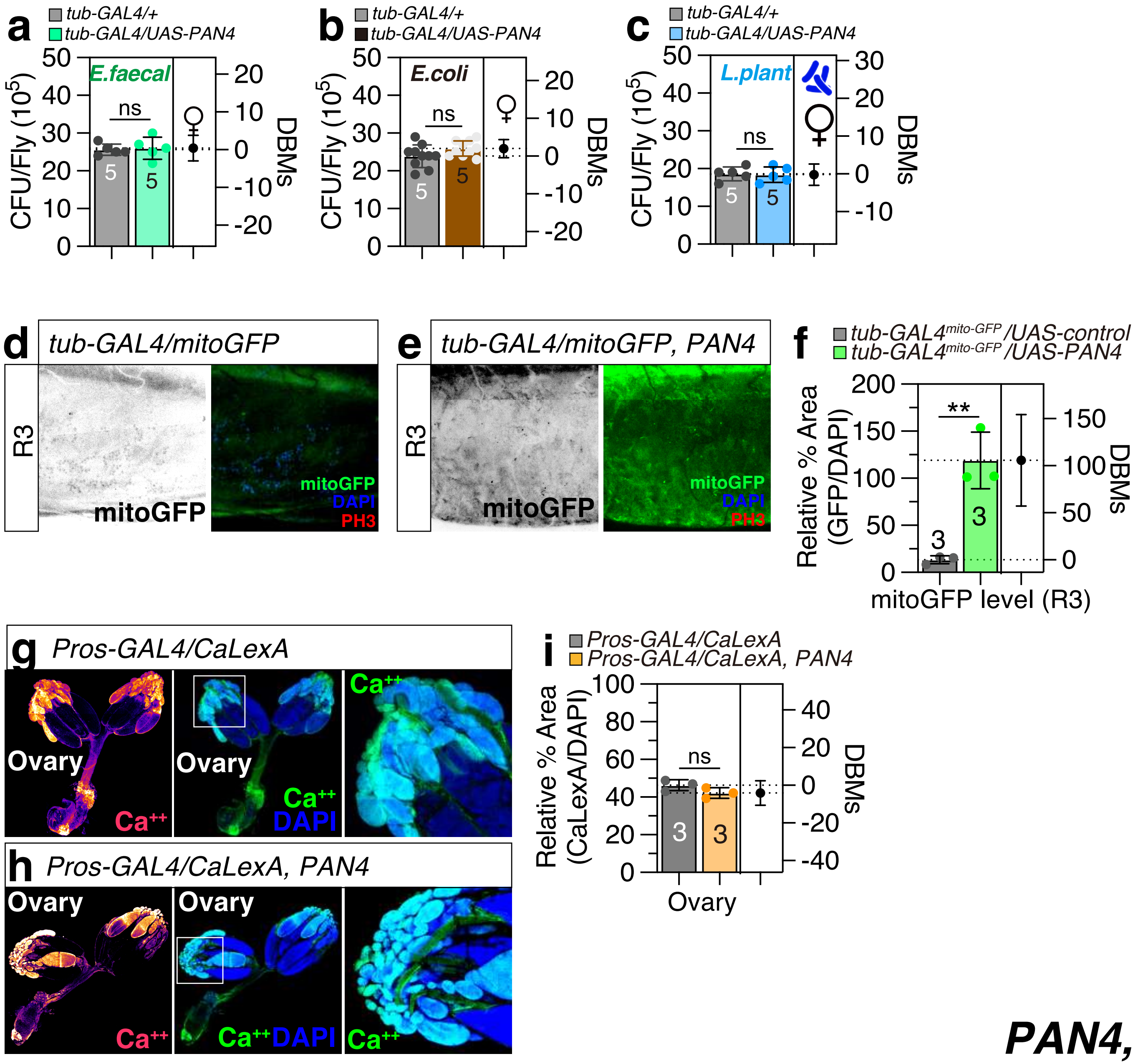

### Supplemental Figure.4

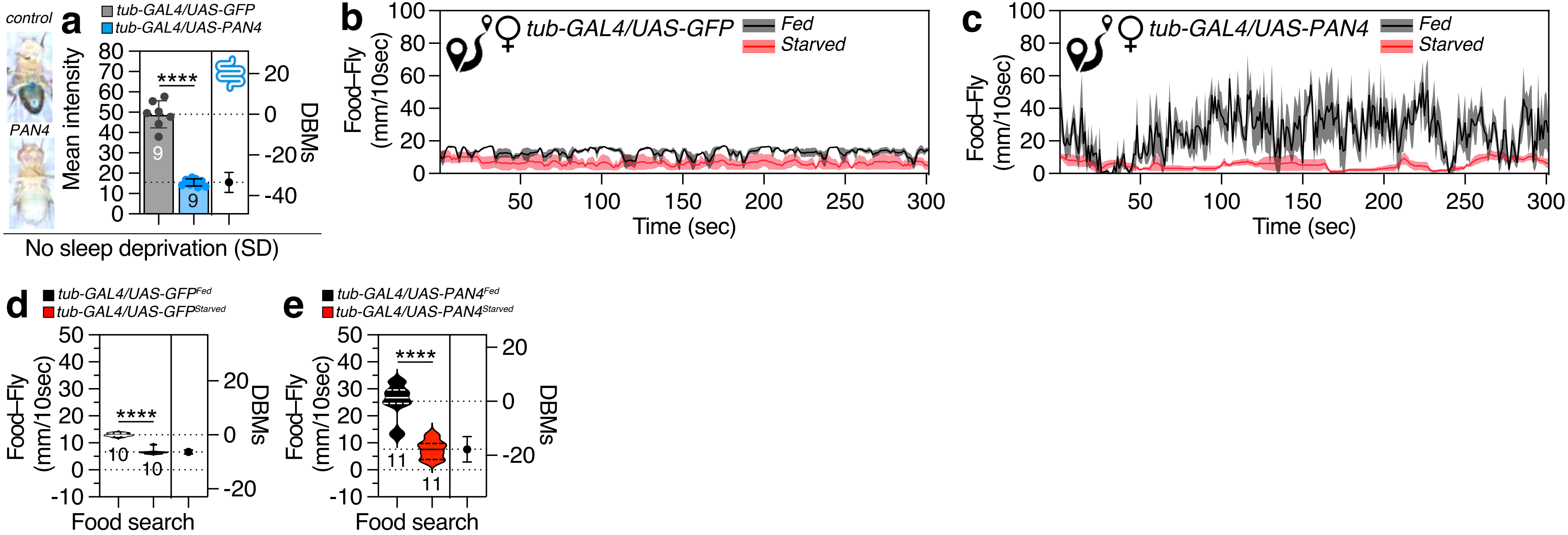
