## Supplemental Figure.2 for "Leveraging *Drosophila* Models to Explore AI-generated Synthetic Peptide’s Potential in Boosting Honeybee Health and Resilience"

**a**

| Name | Sequence | Molecular weight | Net charge at pH7.4 |
|---|---|---|---|
| PAN1 | ALLEKSKK | 916.12 | 1.6 |
| PAN2 | VSLVKAALKE | 1057.28 | 0.52 |
| PAN3 | LLDFKLSDAK | 1149.33 | –0.44 |
| PAN4 | GAYTFKIRRK | 1239.46 | 3.55 |
| PAN5 | FKRAKKWVFR | 1365.67 | 4.55 |

*Surana et al.(2023)*

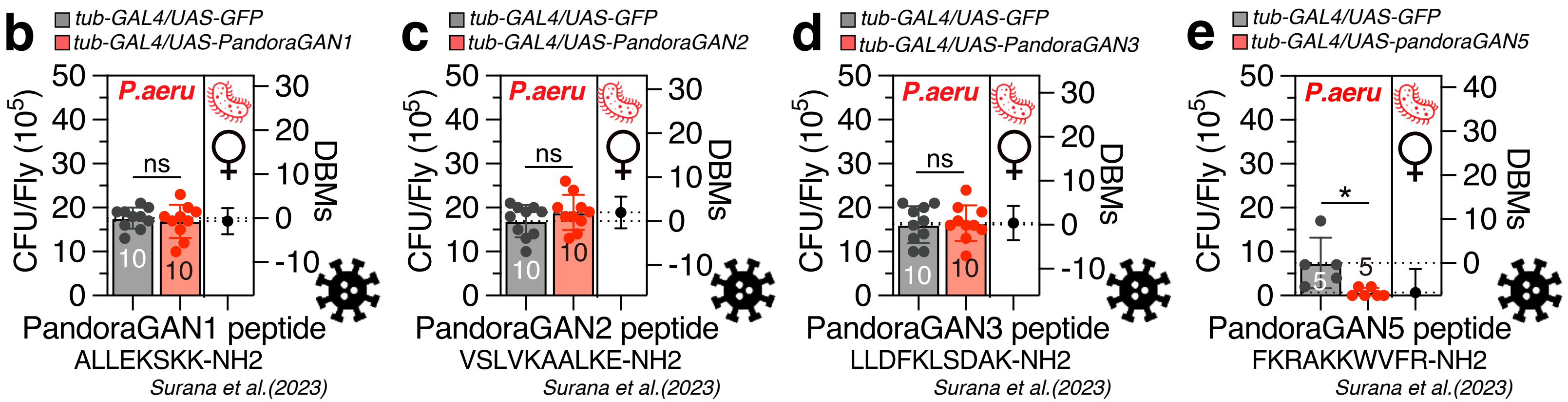


antiviral peptides

***PAN4, Fig.S2***
